# Inertia triggers nonergodicity of fractional Brownian motion

**DOI:** 10.1101/2021.06.17.448813

**Authors:** Andrey G. Cherstvy, Wei Wang, Ralf Metzler, Igor M. Sokolov

## Abstract

How related are the ergodic properties of the over- and underdamped Langevin equations driven by fractional Gaussian noise? We here find that for massive particles performing fractional Brownian motion (FBM) inertial effects not only destroy the stylized fact of the equivalence of the ensemble-averaged mean-squared displacement (MSD) to the time-averaged MSD (TAMSD) of overdamped or massless FBM, but also concurrently dramatically alter the values of the ergodicity breaking parameter (EB). Our theoretical results for the behavior of EB for underdamped ot massive FBM for varying particle mass *m*, Hurst exponent *H*, and trace length *T* are in excellent agreement with the findings of extensive stochastic computer simulations. The current results can be of interest for the experimental community employing various single-particle-tracking techniques and aiming at assessing the degree of nonergodicity for the recorded time series (studying e.g. the behavior of EB versus lag time). To infer FBM as a realizable model of anomalous diffusion for a set single-particle-tracking data when massive particles are being tracked, the EBs from the data should be compared to EBs of massive (rather than massless) FBM.

## I. INTRODUCTION

Fractional Brownian motion (fractional BM or FBM)—introduced by Kolmogorov^1^ and further developed by Mandelbrot and van-Ness^2^—is one of the paradigmatic stochastic anomalous-diffusion processes^3–6^ employed to describe or rationalize numerous experimental observations of non-Brownian or nonlinear diffusion taking place on various length- and time-scales, from the nanoworld to the interstellar space (the list of relevant studies is too long to overview it in this short paper).

In its classical formulation, the dynamics of a single FBM particle is described by the overdamped Langevin equation (the high-friction scenario)^7–13^

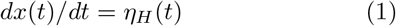

driven by fractional Gaussian noise *η*_*H*_(*t*) with zero mean and power-law long-ranged correlations, with (for *t* ≠ *t*′)

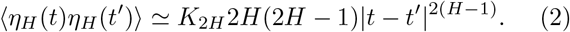

For “athermal” FBM dynamics—with no (generalized) fluctuation-dissipation relation being fulfilled, in contrast to BM—the mean-squared displacement (MSD)

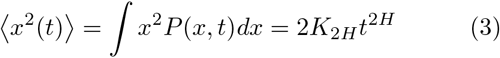

follows from the probability-density function, which for the Dirac-delta-like initial condition, *P* (*t* = 0) = *δ*(*x* − *x*_0_) is given by the Gaussian

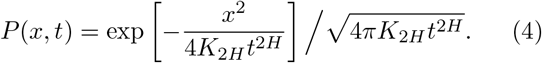

Here, *K*_2*H*_ is the generalized diffusion coefficient (with the physical units [*K*_2*H*_]=m^2^ /sec^2*H*^) and *H* is the Hurst exponent^3,6^. The increments of FBM are positively (negatively) correlated for superdiffusive (subdiffusive) Hurst exponents, for 1 > *H* > 1*/*2 and 0 *< H <* 1*/*2, respectively. Standard BM is FBM at *H* = 1*/*2.

From a time series *x*(*t*) of a stochastic process, the time-averaged MSD (TAMSD) is standardly defined as the sliding average of squared increments along the trajectory of length *T* as

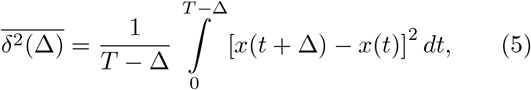

where Δ is the lag time. Averaging over *N* realizations of a random or fluctuating variable 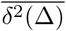 constructed via (5) from *x*(*t*), the mean TAMSD at a given Δ value is

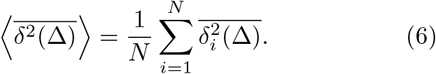

Hereafter, the angular brackets denote ensemble averaging, while time-averaging is shown by the overline. The concept of ergodicity^3,6,15–17^ implies the equivalence of the MSD to the [mean] TAMSD in the limit of long trajectories and short lag times, at Δ*/T* ≪1. FBM is ergodic^7–9,13,18,19^, with

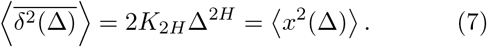

The degree of nonergodicity—or of innate irreproducibility of the TAMSD realizations—is quantified by the ergodicity breaking parameter^6,7,20^,

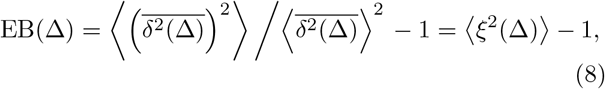

that is the ratio of the variance to the squared mean of 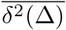. The dispersion and distribution of the TAMSDs (5) around their mean (6) is embodied in the distribution *ϕ*(*ξ*), where 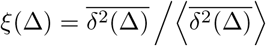.

In the continuous-time framework, EB of FBM for the Hurst exponent 0 *< H <* 3*/*4 deviates only slightly from EB for BM,

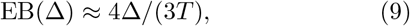

growing at Δ*/T ≪* 1 as^7^ (see Eq. (19) below) EB(Δ) ≈ *C*_1_(*H*) × (Δ*/T*)^1^. For 1 > *H* > 3*/*4 the behavior of EB(Δ) of FBM is more subtle/complicated: the EB values also depend explicitly on the lag time [rather than only on (Δ*/T*)-ratio], with EBs being generally significantly larger. In this range of *H* exponents FBM features a slower approach to ergodicity: we refer to the studies^7,12,14,21–23^ for the continuous- and to Refs.^13,18^ for the discrete-time calculations of EB for FBM.

Our key objective here is to quantify (non-)ergodicity of FBM in terms of EB for diffusion of massive—rather than massless—particles. We focus on the experimentally relevant setup of a constant trace-length *T* and varying particle mass *m*, as often utilized in EB calculations (e.g., for size- or mass-polydisperse tracers in single-particle tracking (SPT) experiments). We use under-damped/massive and overdamped/massless interchangeably for the respective scenarios of FBM diffusion. FBM particles with *m* ≠ 0 require an underdamped description yielding a new time-scale, see below.

The paper is organized as follows. Starting with the description of the simulation scheme in Sec. II A, in Sec. II B the results for the MSD and mean TAMSD of under-damped FBM are presented, both from computer simulations and theory. In Sec. II C the main results for the EB parameter of massive FBM are presented and contrasted to those for massless FBM. We conclude in Sec. III and mention some applications. In App. A certain technical details and derivations are given, while some auxiliary figures are presented in App. B.

## II. MAIN RESULTS

### A. Diffusion model and simulation scheme

We simulate the diffusion for a single tracer of mass *m* driven by external fractional Gaussian noise *η*_*H*_ (*t*) using the underdamped Langevin equation,

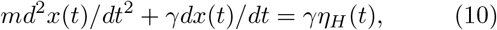

used recently also in Refs.^24–27^. Here, *γ* is the friction coefficient, that is set to *γ* = 1 below, so that the characteristic momentum-relaxation or Smoluchowski’s time,

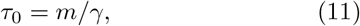

is controlled solely by particle mass *m*. To simulate (10) we introduce the velocity *v*(*t*) = *dx*(*t*)*/dt* and discretize the resulting equations (A3) with the time-step *dt* to get

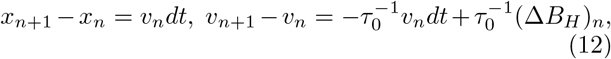

with 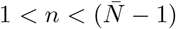,where

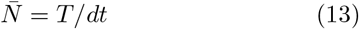

is the number of elementary steps in the trajectory. This system of forward recurrent equations is solved with the initial position *x*_0_ and velocity *v*_0_ of the particle. Both *v*_0_ = 0 and *v*_0_ being distributed according to the stationary-state distribution are considered. The increment of standard FBM (Δ*B*_*H*_)_*n*_ are generated using the Wood-Chan method [faster than the Hosking approach] based on the fast-Fourier transform^28^.

### B. MSD and TAMSD

In Fig. 1 we present the results of simulations for the MSD and TAMSD of massive-FBM particles starting their motion with *v*_0_ = 0, for two values of the Hurst exponent *H* (for sub- and superdiffusive dynamics). We find that the long-time dynamics of massive FBM is expectedly the same as for massless FBM, with the MSD (3) being equal to the TAMSD (7). At short lag times the mean TAMSD of massive FBM start ballistically,

**FIG. 1:**
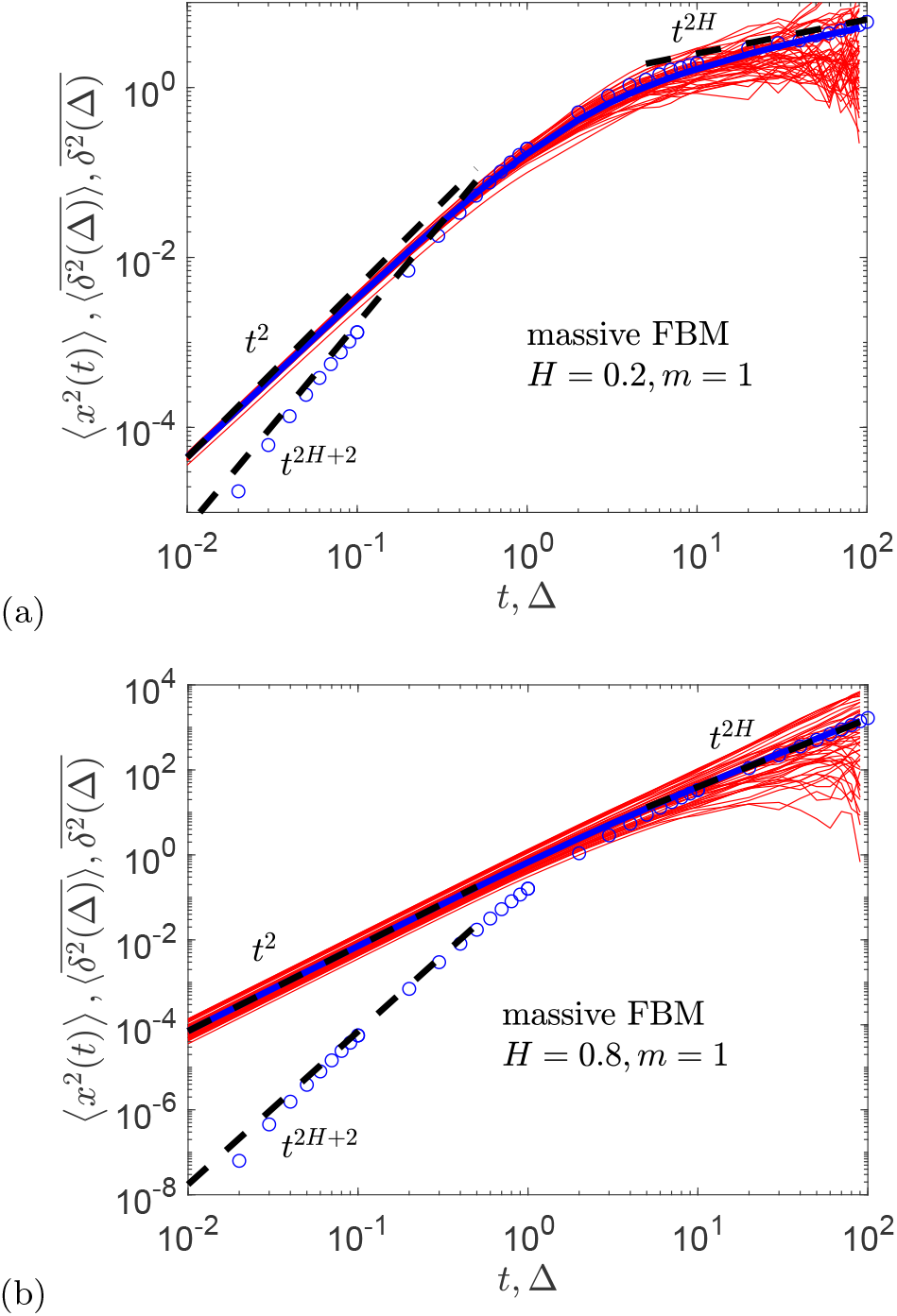
Magnitude of the MSD ⟨*x*^2^(*t*) ⟩ (blue circles), the spread of individual TAMSDs 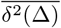 (thin red curves), and the mean TAMSD 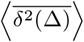 (thick blue curve) for massive FBM, shown for two values of the Hurst exponent (see the legends). The long-time MSD and mean TAMSD asymptotes given by expressions (3) and (7), respectively, are the dashed lines. The short-time asymptotes (15) and (14) for the MSD and mean TAMSD are the dashed lines. Parameters: the particle mass is *m* = 1, the time-step of simulations is *dt* = 10^−2^, the trajectory length is *T* = 10^2^ [or 10^4^ time-steps], the number of independent trajectories used for ensemble averaging is *N* = 10^3^, the initial conditions are *x*_0_ = *v*_0_ = 0, and the generalized diffusivity is set at *K*_2*H*_ = 1*/*2.

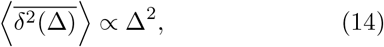

while its MSD grows super-ballistically,

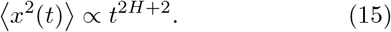

The exact prefactors in these relations are derived in Apps. A 1 and A 2, see Eqs. (A28), (A30) and (A19), respectively. In terms of the MSD-to-TAMSD equivalence, underdamped FBM ceases to stay ergodic: both scalings and magnitudes of ⟨*x*^2^(Δ) ⟩ and 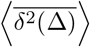 are disparate, see Fig. 1. Massive FBM also features larger EB values, see Sec. II C.

The long-time evolution of the MSD and TAMSD for massive FBM is identical, given by that of massless FBM in (3) and (7), namely

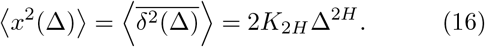

The temporal extent of the new short-time regimes given by (14) and (15) grows with the particle mass (similarly to that for massive BM described by the position-Ornstein-Uhlenbeck (OU) process^29^) and depends on the Hurst exponent. These two parameters determine the “agility” of the underlying dynamics, compare Figs. 1 and S1 for the 10-times heavier particles. Expressions (A22) and (A34) give the crossover times to the long-time behavior (16) of the MSD and mean TAMSD.

We emphasize that initial velocities of the particles— instead of being zero as in Fig. 1—can be sampled from the stationary distribution (A12), with ⟨ *v*^2^ ⟩ _st_ given by (A11). Then, the MSD and mean TAMSD are *equal* and both acquire a transient short-time ballistic growth,

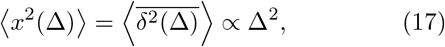

with a time of transition to the long-time behavior given by expression (A23). For stationary-state-distributed initial velocities, the process of massive FBM stays “ergodic” at short times in the sense of the MSD-to-TAMSD equivalence, whereas the long-time growth of the two averages is the same as for massless FBM given by expression (16), see Fig. S2 as well as Apps. A 1 and A 2. We stress, however, that despite (17) is true for the averages, the irreproducibility of individual TAMSDs increases for larger Hurst exponents, as quantified in Fig. S2b. This increase has dramatic implications for the EB values characterizing the dispersion of 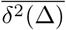 around 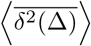, see Sec. II C.

Analogously to equipartition theorem, at long times we can define the effective *H*-dependent “temperature”,

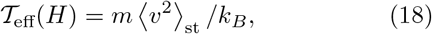

where *k*_*B*_ is the Boltzmann constant. For our nonequilibrium system the width of the Gaussian velocity distribution in the stationary state given by (A12) and the mean-squared velocity of the particles ⟨ *v*^2^ ⟩ _st_ given by (A11) are thus controlled by 𝒯_eff_ (*H*).

### C. EB

#### 1. Spread of 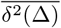

From Fig. 1 it becomes apparent that for superdiffusive underdamped FBM the individual TAMSDs become less reproducible. This fact gets reflected in larger values of the EB parameter, as compared to those for standard FBM^7,13^, see the results of simulations in Fig. S3 for EB(Δ) for FBM with *H* = 0.8 and 0.2 (both for the under- and overdamped cases). The increase of EB for massive FBM is observed in the entire range of lag times, but at the shortest lag time Δ = Δ_1_ = *dt* it is particularly dramatic. This is associated with a larger degree of nonergodicity (in terms of EB^6^). This fact is relevant for SPT-experiments as—in virtue of the best time-averaging statistics at short lag time—the time-series are often used to compare the assessed EB(Δ_1_) values to EBs of known models of diffusion^6,30–32^. We stress that for the massive particles driven by *η*_*H*_ —with the initial conditions *x*_0_ = *v*_0_ = 0—the EB values are dramatically larger than those for overdamped FBM^7^. The deviations of massive-versus massless-FBM results for EB are (expectedly) the largest for lag times where the respective MSD and TAMSD deviate, compare the respective Δ-regions in Figs. 1 and S3. The number of independent *in-silico*-generated trajectories used for ensemble averaging in Fig. 1 and all later plots is at least *N* = 10^3^.

#### 2. EB of massless FBM

In Fig. S3 we show for comparison the theoretical predictions for EB of massless FBM in the framework of *continuous* time given by^7,12,13^

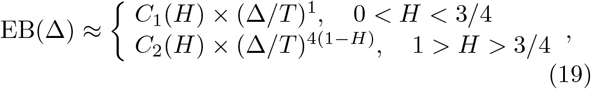

with the coefficients

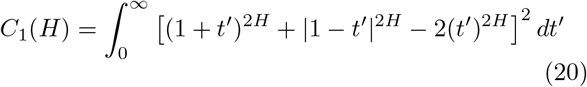

(see Fig. 8 of Ref.^13^ for *C*_1_(*H*) values for some *H*) and

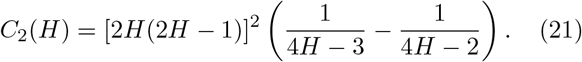

Here, (Δ*/T*) is the only possible dimensionless time.

We also use the results for EB of massless FBM in the *discrete*-time approach^13,22^, predicting that at the first step is (for *H <* 1*/*2)

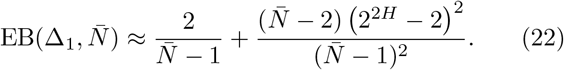

Here, 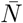 is given by Eq. (13). For *H* > 1*/*2 the result for 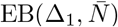) is more complicated, given by Eq. (C4) of Ref.^13^. The terminal value (see Ref.^13^ for details)

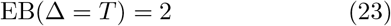

is also referred to in Fig. S3 and in other EB plots.

In Fig. S4 the variation of EB(Δ) of massive FBM—computed at the shortest lag time Δ = Δ_1_—with the Hurst exponent *H* is summarized. For underdamped FBM the behavior of EB(Δ_1_) is very different from the discrete-time prediction (22) (see our recent study^13^ for detailed derivations and description). Instead, we find that, starting from very small *H*, the values EB(Δ_1_) grows quickly with *H*, rather than stay nearly constant in the range 0.6 ≳ *H* > 0 as EB expression (22) would predict. At Δ → *T* the value of EB(Δ) approaches (23).

#### 3. EB of massive FBM

For a fixed *m*, for longer times series massive FBMs turn progressively “less heterogeneous” and deviations from ergodicity decrease. Specifically, EB(Δ_1_, *T*) values obtained from simulations are found to decrease with the trajectory length *T* according to the empirical scaling

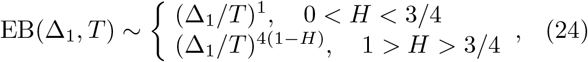

as illustrated in Fig. 2. The approach to ergodicity for longer trajectories of massive-FBM particles is, thus, the same as that for standard FBM^7^ in expression (19), with the anomalously slow EB decrease with *T* and slower approach to ergodicity for 1 > *H* > 3*/*4. As follows from Fig. 2, for a fixed mass *m*, a slower decay of the EB parameter for massive FBM for increasing *H* requires longer trajectories for scaling (24) to be applicable. For comparison, the empty symbols in Fig. 2 are the results of simulations for the decay of EB(Δ_1_, *T*) with *T* for massless FBM following Eq. (19).

**FIG. 2:**
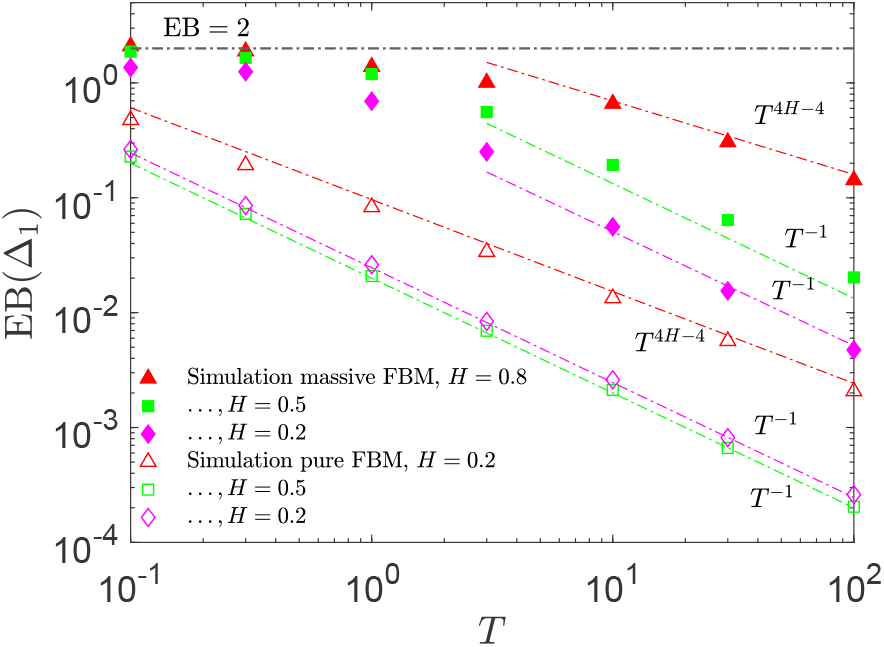
EB of massive FBM versus the trajectory length *T* for several values of the *H* exponent (the filled symbols), with the scaling relations (24) and (A50) shown as the dashed lines [no data fitting here, the exact prefactors used]. The results of EB simulations of massless FBM for the same *H* are the empty symbols; they are shown together with the modified Deng-Barkai^7^ EB asymptote (19) and discrete-time EB expression (21), see Ref.^13^. Parameters: *m* = 1, *dt* = 10^−2^.

Our main results, presented in Fig. 3, show that for vanishing mass *m* the EB values for massive FBM approach those of massless FBM, Eq. (22) in the discrete-time approach, as expected. For extremely massive particles, the EB parameter reaches its maximal values, saturating at *H*-dependent plateaus (see Eq. (A40) and the details of the derivations in App. A 3), and also Fig. S5. These large EB values corroborate with the most pronounced spread of 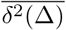 and, therefore, broader distributions *ϕ*(*ξ*(Δ_1_)) for diffusion of heavier FBM particles, as detailed in Fig. S1 (to be compared to Fig. 1).

**FIG. 3:**
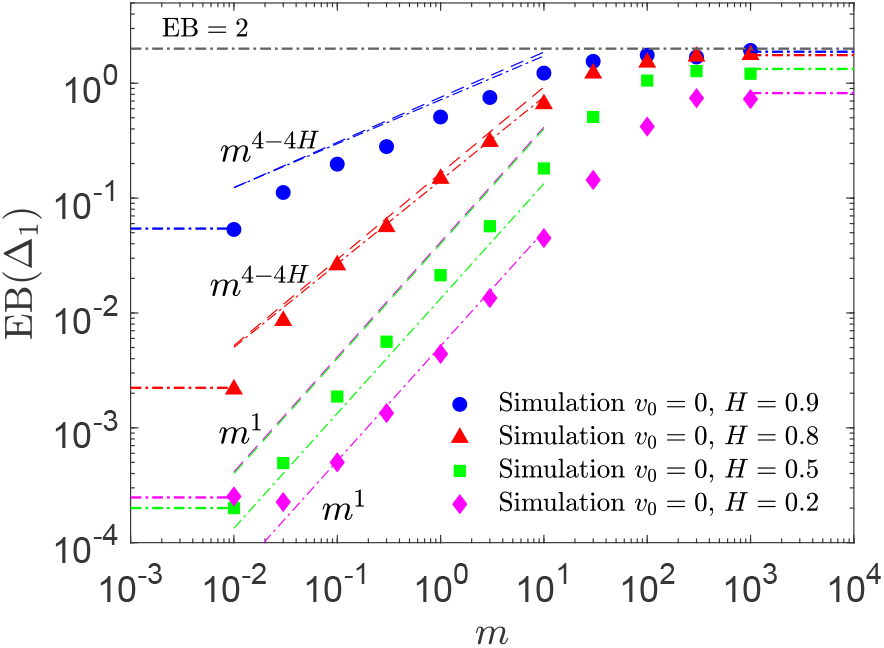
EB parameter of underdamped FBM plotted versus the particle mass *m* for several *H* values. At intermediate *m*-values we show the scaling relations (25) and (A48) as the dashed colored lines and the EB expression (A50) with the exact prefactors [no data fitting] as the dot-dashed colored asymptotes. The terminal EB value (23) is also shown. The colored dot-dashed lines on the left and on the right side of the plot, respectively, indicate the EB discreteness-induced plateaus for massless FBM in the discrete-time formulation given by expression (22) and the EB plateaus in the large-mass limit given by (A40). Parameters: *T* = 10^2^, *dt* = 10^−2^.

For the particles of intermediate masses, we empirically find scaling relations

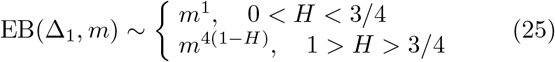

which are in excellent agreement with the *in-silico* results for all *H* values, see Fig. 3. In App. A 3 we present the derivations of the EB plateaus for large-mass or short-trace conditions given by (A40) and the variation of EB for long trajectories given by expression (A48), both in excellent agreement with the results of simulations and identical to the empirical laws (24) and (25).

We stress that, although for the case of stationary-state-distribution of initial velocities (A12) of the massive-FBM particles the TAMSD scaling does get affected and the short-time MSD-versus-TAMSD equivalence gets restored, as shown in Fig. S2, using the stationary-state distribution for *v*_0_ *does not change* the scaling relations for EB(Δ_1_, *m*) versus mass *m* as given by expression (25), affecting only the EB-plateau values at *m* → ∞, see Fig. S6.

## III. DISCUSSION AND CONCLUSIONS

We examined the diffusion process driven by fractional Gaussian noise *η*_*H*_ in the presence of inertia, both analytically and via stochastic computer simulations. We found that the short-time transient mass-dependent growth of the MSD is superballistic with ⟨ *x*^2^(*t*) ⟩ ∝ *t*^2*H*+2^, while the mean TAMSD grows at short-lag-time ballistically, 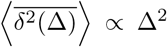. In this range of times—quantified in Apps. A 1 and A 2—a pronounced inertia-induced MSD- to-TAMSD nonequivalence was found. Systematically varying the Hurst exponent *H* and particle mass *m* we demonstrated that the evolution of the EB parameter at short lag times follows the novel scaling relations with trajectory length *T* and tracer mass *m*. In particular, EB for massive FBM grows with *m* at a fixed *T*, reaching much larger values than EBs of massless FBM. The scaling relations (24) and (25)—derived in App. A 3 based on solid theoretical grounds as (A48) and (A50)—are closely supported by the results of simulations and embody the main findings of this study.

We stress that the effects of inertia, inevitable in the short-time behavior of real-world tracers in SPT-experiments, should be carefully accounted for if a quantitative comparison of the computed EB values versus those for FBM (or other diffusion models) is to be performed. For short lag times—where the EB-averaging statistics is most accurate and, thus, often used in the SPT-data analysis—the values of EB were found to be most affected by these inertia effects. The emerging “ballisticity” in the TAMSD and the altered scaling of the MSD in this short-Δ region was shown to trigger the MSD-to-TAMSD nonequivalence and weak ergodicity breaking for the *η*_*H*_-driven dynamics of massive particles.

From the experimental perspective, the *m*- and *H*-dependent time-scales—when the above-mentioned ballistic effects can play a role in the massive-FBM dynamics (see Eqs. (A22) and (A34))—should be understood for a given tracer mass *m*, trajectory length *T*, lag time Δ, and properties of the diffusion medium *γ*. This will clarify whether the short- or long-time asymptotic behaviors of the TAMSD and EB are valid for a given set of data and parameters. The computed EB values—if aimed at a quantitative comparison with known models of massless-particle diffusion—should be properly *adjusted* to account for a finite particle mass, *m*. This is exemplified above for massive FBM.

From the theoretical perspective, the velocity-correlation time *τ*_0_ = *m/γ* naturally rescales the time-scale *T* to be taken for a physically correct comparison of EB values for FBM computed for the particles of varying mass. As we have shown, the inertial effects make FBM appear nonergodic when, e.g., the trajectory length *T* is fixed while mass *m* is varied. A correct rescaling of the trace-length *T* with a typical relaxation time *τ*_0_ would be

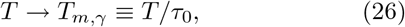

so that heavier particles diffuse for the same *relative* time, *T*_*m,γ*_. The relaxation time *τ*_0_ is, therefore, the new physical time-scale, as compared to a “scale-free” massless FBM. The mass-dependent *T* -rescaling (26) renders the respective EB(Δ = Δ_1_) values *universal and constant* for a fixed *T*_*m,γ*_, i.e.

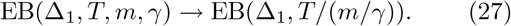

We have indeed confirmed this invariance of rescaled EBs via simulating massive FBM for a constant *T*_*m,γ*_ but for a set of different *T* and *m* values, see Fig. 4, for both sub-and superdiffusive *H* exponents. Note that a plateau-like behavior of EB at small Δ (especially for *H* = 0.8), as shown in Fig. S3, enables us to consider in Fig. 4 for rescaling (27) the EB values at the same Δ_1_ for different *T*. The rescaling (26) is also supported by theoretical arguments in App. A 3: namely, EB in (A48) varies with the rescaled (*τ*_0_*/T*)-variable only. We stress that a similar time-rescaling “regularizes” apparent nonergodicity emerging for FBM with stochastic resetting^18,19^.

**FIG. 4:**
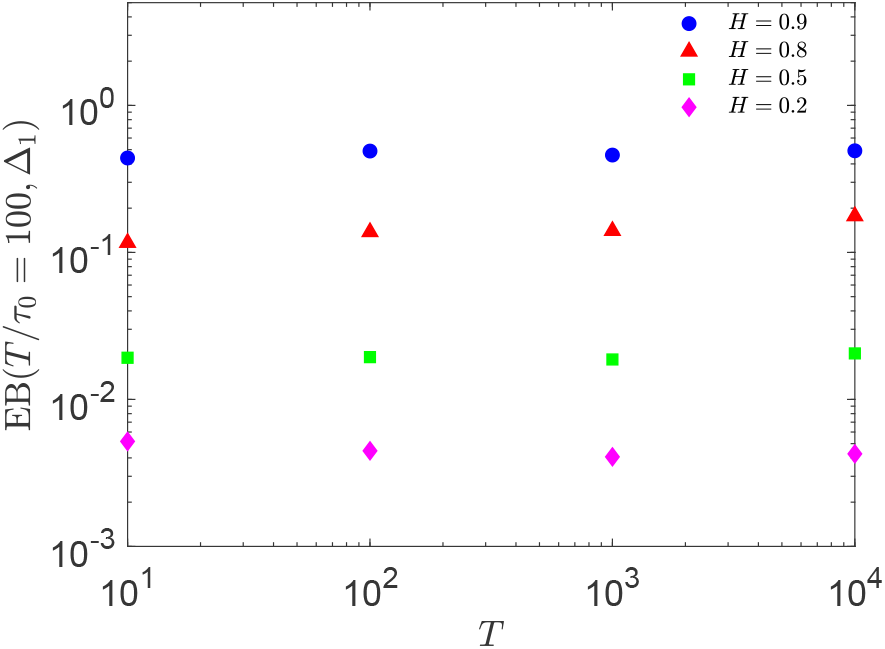
EB of massive FBM at Δ_1_ = 10^−2^ for *simultaneously* varying particle mass *m* and trajectory length *T* such that the *rescaled* trace-length (26) stays constant, namely *T /τ*_0_ = 10^2^.

Note that the theoretically desirable rescaling (26) might not always be realizable in SPT experiments, performed, e.g., for an ensemble of mass-polydisperse tracers. Certain restrictions of measurement protocols and limitations of the apparatus itself [in terms of recording time, measurement precision, etc.] can prevent such an idealized scenario from happening. As a result, non-rescaled as per (27) values of the EB parameter will be extracted from the SPT time-series. We therefore presented the main theoretical results for variation of the EB parameter of massive FBM and pronounced emerging nonergodicity in such a practice-related setup.

The nonergodicity of underdamped FBM is therefore “apparent”, being the direct consequence of insufficient trajectory lengths for progressively heavier particles [the time, required for a system to visit cells in phase-space in the same proportion to attain a similar degree of ergodicity]. This is similar to the “apparent” nonergodicity predicted, for example, in a two-state switching-diffusion model^33^ [with, respectively, {*k*_1_, *D*_1_} and {*k*_2_, *D*_2_} being the switching rates and diffusivities]. Such a model (see also Refs.^34,35^) becomes “apparently” profoundly nonergodic when the mean resident time in a given diffusion state exceeds the overall length of the trajectory (i.e., for occupation times^33^ *τ*_1,2_ = 1*/k*_1,2_ ≳ *T*).

Note here also that above we followed the definitions of the MSD and TAMSD standard for the SPT-data analysis, although some alternative “next-level” definitions of (non-)ergodicity can [in some situations] be theoretically more appropriate for assessing the MSD-vs-TAMSD equivalence (see, e.g., Refs.^36,37^).

Finally, this analysis continues a series of recent studies of (non-)ergodic dynamics of massive particles for anomalous-diffusion processes with a power-law *D*(*t*)^26,27,38^ and exp-log form of *D*(*t*)^39^. The extension of the “nonzero-mass concept” for unveiling the MSD-vs-TAMSD nonequivalence and EB-based nonergodicity is of interest also for other models of anomalous diffusion^4,6,40,41^, both for pure and “modified” processes [e.g., by external confinement^9,42^, TAMSD-ageing effects^27,43^, in barrier-escape setups^44^, and for stochastic-resetting protocols^18^].

## Acknowledgments

A. G. C. gratefully acknowledges the Humboldt University of Berlin for hospitality and support. R. M. acknowledges financial support by Deutsche Forschungsgemeinschaft (DFG Grant ME 1535/12-1). R. M. thanks the Foundation for Polish Science (Fundacja na rzecz Nauki Polskiej) for support within an Alexander von Humboldt Polish Honorary Research Scholarship.

## Abbreviations

BM: Brownian motion
FBM: fractional BM
MSD: mean-squared displacement
TAMSD: time-averaged MSD
SPT: single-particle tracking
OU: Ornstein-Uhlenbeck

## Appendix A: Derivation of the MSD, mean TAMSD, and EB for massive FBM

### 1. MSD

#### a. General solution

The Langevin-type equation of a single massive-FBM particle of mass *m* can be written as

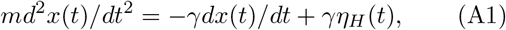

where *γ* is the damping coefficient. In the overdamped limit, the inertia term in Eq. (A1) can be neglected yielding the massless-FBM formulation^2,7–9,13^. The external-noise intensity, scaling as 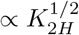 as per Eq. (2), is generally not coupled to the friction coefficient. With the particle velocity

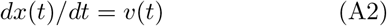

Eq. (A1) is the velocity-OU process^29^ for *v*(*t*) driven by fractional Gaussian noise *η*_*H*_ with the correlator (2), i.e.

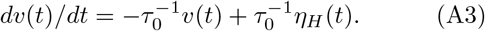

Here, *τ*_0_ = *m/γ* is the velocity-relaxation time (11). The formal solution of Eq. (A3) with the initial condition *v*(*t* = 0) = *v*_0_ is

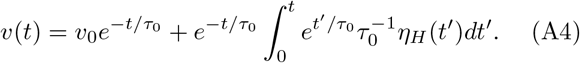

The noise-averaged squared velocity is given by^37^

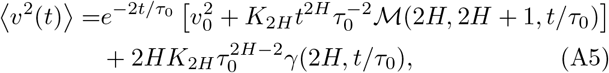

where *B*_*H*_ (*t*) is FBM, ℳ (*a, b, z*) is the confluent hyper-geometric (or the Kummer’s) function of the first kind,

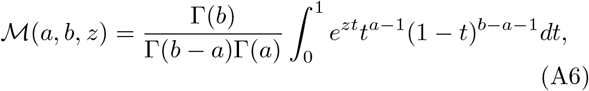

and *γ*(*a, z*) is the lower incomplete Gamma function,

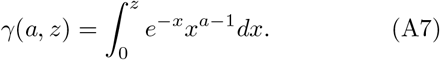

The expression (A5) follows from Eq. (28) of Ref.^37^ [see also the detailed derivation there] after setting the relaxation time to *τ*_0_ and the actual noise-strength as in Eq. (A3), after some algebra with the ℳ-functions^46^.

#### b. Short- and long-time limits

At short times, when

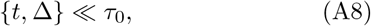

using the small-argument expansions ℳ (2*H*, 2*H* + 1, *t/τ*_0_) ≈ 1 and *γ*(2*H, t/τ*_0_) ≈ (2*H*)^−1^ (*t/τ*_0_)^2*H*^, one gets from (A5) that

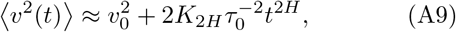

whereas at long times, at

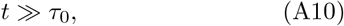

using the respective expansions 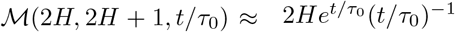 and *γ*(2*H, t/τ*_0_) ≈ Γ(2*H*), where Γ(*x*) is the Gamma function, we find stationary, time-independent velocities of the particles in the parabolic OU potential. The long-time, stationary-state-related mean-squared velocity ⟨ *v*^2^(*t*) ⟩ follows then from (A5) as

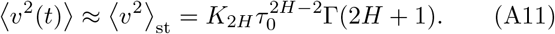

The Gaussian distribution of long-time velocities,

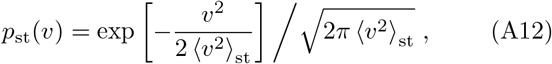

is akin to the 1D Maxwell-Boltzmann distribution with 𝒯_eff_ defined via (18). The distribution (A12) satisfies the stationary Fokker-Planck equation for the fractional velocity-OU process (A3) given by^47,48^

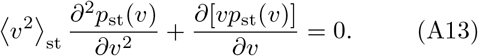

The asymptotic result for the velocity-autocorrelation function of the fractional OU process for the stationary velocities for the well-separated time instances, at *δt* ≫ *τ*_0_, is given by^37^ [see also Ref.^49^]

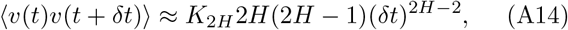

while at *δt* ≪ *τ*_0_ we have

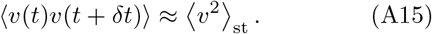

As *m, γ*, and noise strength are independent model parameters, to get simpler solutions and to capture main physical scalings, we consider now the “large-mass”- or “short-trajectory”-limit of Eq. (A3) via setting *τ*_0_ to a large value, but such that the product 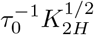 stays constant via adjusting the noise-strength [neglecting, thus, the second term in (A3)]. The evolution of velocity at short times, starting from *v*_0_ value, is then simply integrated *η*_*H*_ (*t*) that is FBM, namely

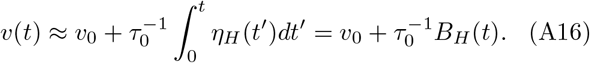

The velocity autocorrelation function and second moment are then given, respectively, by

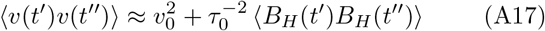

and

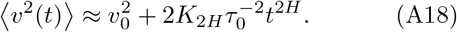

As per Eq. (A2), the particle position *x*(*t*) is one more integration of (A16), the MSD in this limit for the initial condition *x*(*t* = 0) = 0 is a combination of a ballistic ∼ *t*^2^and a faster-than-ballistic ∼ *t*^2*H*+2^ term, namely

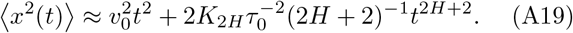

#### c. Distributed v_0_

Assuming initial velocities to be distributed with the stationary-state-distribution (A12), the MSD at short times—after neglecting the second term in (A19)—reads

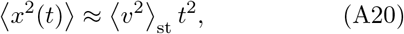

while at long times (A10) [neglecting the smaller-in-magnitude first term in Eq. (A19)] one gets

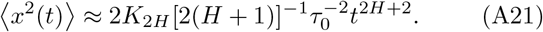

We note that the long-time MSD behavior is observed beyond the transition time 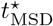 scaling as

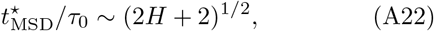

and

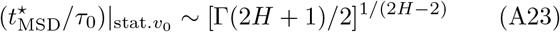

for the case *v*_0_ = 0 and *p*_st_(*v*)-distributed initial velocities of the particles, correspondingly. For a [vanishingly] small mass, neglecting the first term in (A3), one gets the expected MSD of massless FBM,

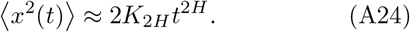

### 2. TAMSD

#### a. Short-time limit

For the TAMSD (5), formally integrating expression (A2) from *t* to *t* + Δ, we get

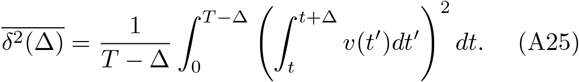

For the region of short lag times, when (A8) holds, neglecting the variation of velocity *v*(*t*) on the time-scale of Δ, we find from (A25) that

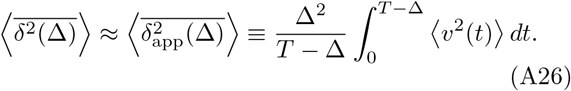

For short trajectories or large particle mass, i.e. at

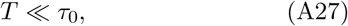

when velocities did not yet reach stationarity, after using the approximate velocity (A9) the TAMSD form (A26) yields—similarly to the MSD representation (A19)—a combination of two terms,

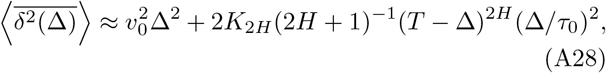

both featuring a ballistic TAMSD growth at *T* ≫ Δ.

#### b. Long-time limit

For long trajectories, when

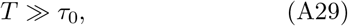

and, thus, for nearly stationary particle velocities, we split the integral in (A26) into two regions, with 0 *< t < τ*_0_ and *τ*_0_ *< t <* (*T* − Δ). In the first region we use the short-time expansion of ⟨*v*^2^(*t*) ⟩ near 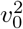 given by (A18), while in the other region the long-time result (A11) with ⟨*v*^2^(*t*) ⟩≈ ⟨ *v*^2^ ⟩ _st_ is used. With this strategy, after integration, the approximate mean TAMSD in this limit becomes

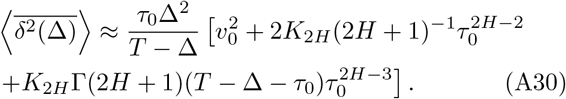

From this expression, neglecting the duration of Δ and *τ*_0_ compared to the trace length *T*, one gets the expected short-lag-time ballistic scaling,

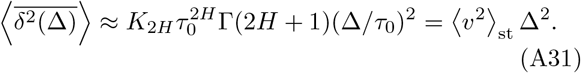

This asymptotic TAMSD is identical to the MSD evolution under the same conditions, Eq. (A21). Therefore, the ergodicity in terms of MSD-to-TAMSD equivalence at short lag times is restored of long trajectories and thus stationary-state velocities.

When both the measurement time and the lag time are longer than the velocity-relaxation time-scale, i.e. at *τ*_0_ ≪ {Δ, *T*}, after using the correlator

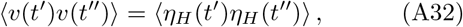

following from Eq. (1) with the initial velocities being fully relaxed by that long time, for the mean TAMSD— starting from expression (A25) and performing the elementary integration—we find

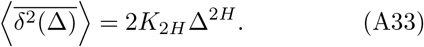

The MSD-to-TAMSD equivalence is thus again restored. The transition lag time from the short-time TAMSD behavior (A31) to the long-time asymptotic law (A33), denoted as 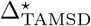, is the same as that for the MSD case with stationary-state-distributed-*v*_0_ given by (A23),

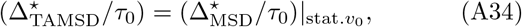

### 3. EB

The standard procedure for computing the EB parameter is to find the general expression for the 4th-order correlation functions of particle positions [for Gaussian random variables] and with its help express all nine terms in the integrand of the 4th-moment 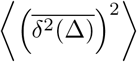 in EB in Eq. (8). This general methodology is often tedious [we refer to EB of the OU process computed like this in Ref.^45^]. Our goal here is—via avoiding such exact but lengthy derivations that are not easy to use in practice [i.a., because of special functions often involved which demand a considerable time to plot and to find relevant scaling exponents]—to employ an approximate EB-derivation to get the main physically relevant scalings for EB of massive FBM.

#### a. Short-time limit

To pursue this goal, for the region of short lag times defined by (A8), using the approximate TAMSD 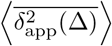 in (A26) and employing Isserlis-Wick theorem for zero-mean Gaussian processes,

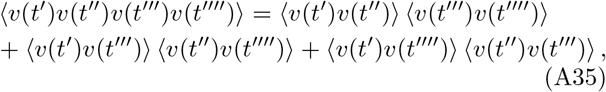

for computing the 4th-order correlators in terms of the pair correlators in Eq. (8), the EB parameter of massive FBM for short trajectories or large particle mass can be approximated as

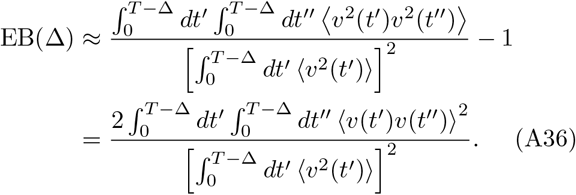

Using the velocity-velocity correlator (A17), the FBM autocorrelation function

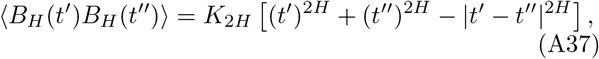

and setting *v*_0_ = 0, for short-*T* or large-*m* values (when the condition (A27) holds) the integrand in the nominator of expression (A36) becomes

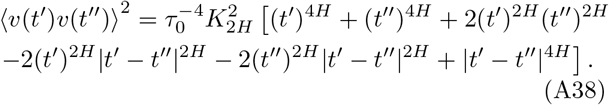

With this result, taking the elementary integrals in (A36), using the symmetry with respect to swapping *t ′* ↔ *t ″* in the integrals, and employing that

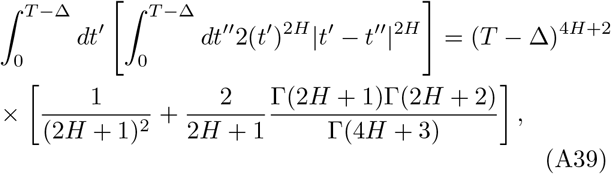

the EB parameter in this limit has no dependence on the lag time and attains the *H*-dependent value

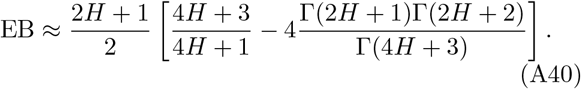

The values of EB in expression (A40) for a set of typical Hurst exponents 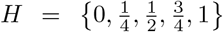 are as follows 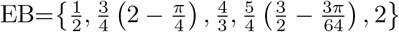. The results of computer simulations for large particle mass for zero initial velocities (the EB values on the right side of Fig. 3) are demonstrated to be in quantitative and excellent agreement with the theoretical prediction (A40), Fig. S5.

For initial velocities *v*_0_ distributed with the stationary-state-distribution, as per (A12), the same EB-computation strategy in the short-time- or large-mass-limit can be employed. Specifically, we start with the general EB form (A36) and use the second moment (A18) and the autocorrelation function (A17) with 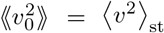 given by (A11). The double angular brackets denote averaging over realizations of initial velocities. After integration, the nominator of EB in Eq. (A36) becomes

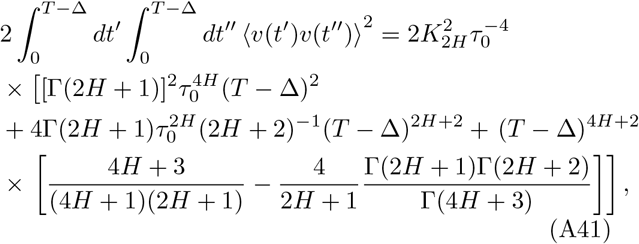

while the denominator is given by

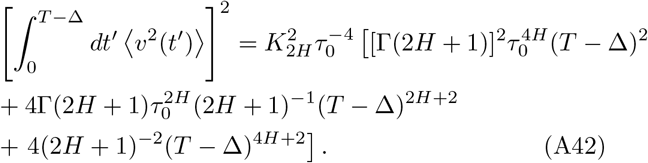

In the limit of large mass or short traces, at *T/τ*_0_ ≪ 1, the highest powers of *τ*_0_ in these expressions dominate yielding in the leading order

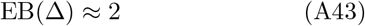

for all values of the Hurst exponent *H*. The value (A43) is shown as the large-mass EB plateau in Fig. S6 and several other EB plots.

#### b. Long-time limit

At long measurement times, when condition (A29) is valid, the velocities executing the fractional OU process are stationary, ⟨*v*^2^(*t*) ⟩ → ⟨ *v*^2^ ⟩_st_. Starting from the representation (A36), the leading-order approximation of EB is

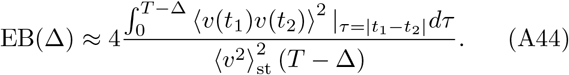

The velocity-autocorrelation features different forms for *τ* ≪ *τ*_0_ given by (A14) and for *τ* ≫ *τ*_0_ in Eq. (A15). Similarly to the approximate TAMSD evaluation in the long-time limit in (A30), we split the integral in expression (A44) into two parts, utilize these different velocity-velocity correlators, and get

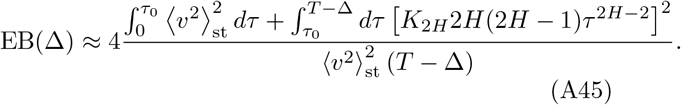

After elementary integration, neglecting the terms of order Δ*/T* ≪ 1, the value of EB becomes independent on the lag time, obeying the relation

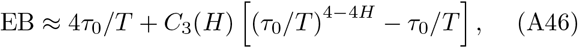

where the coefficient *C*_3_ reads (see Fig. S7 showing its variation)

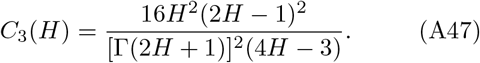

In EB expression (A46), the lag time Δ is effectively replaced by the characteristic time-scale of the problem, *τ*_0_. In the limit *τ*_0_*/T* ≪ 1, after retaining the leading-order terms in (A46), one gets [similarly to the two-region solution for EB of massless FBM given by (24)] the following scaling relations

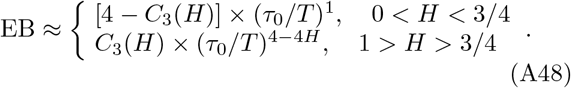

As *τ*_0_ = *m/γ*, the relations (24) and (25) in the main text—used to rationalize the simulation data for EB(Δ_1_, *T, m*) obtained for varying trace length *T* and particle mass *m*—instantly follow from (A48).

As a next-order approximation, instead of (A15) we use the small-increment-expansion in the stationary regime

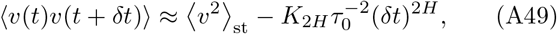

while for the well-separated increments expression (A14) is still employed. Then, repeating the calculations starting from the EB expression (A45), one gets the same scalings as in (A48), but with the improved coefficient for the case 0 *< H <* 3*/*4, namely

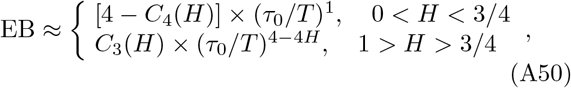

where

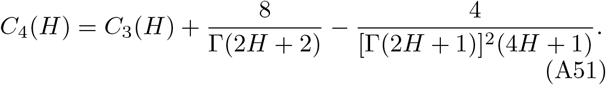

The variation of coefficients *C*_3_(*H*) and *C*_4_(*H*) with the Hurst exponent *H* is shown for completeness in Fig. S7. Using (A50) enables a *quantitative description* of the EB-versus-*m* data extracted from simulations, see Fig. 3.

## Appendix B: Supplementary figures

Here, we present some auxiliary figures supporting the claims of the main text.

**FIG. S1:**
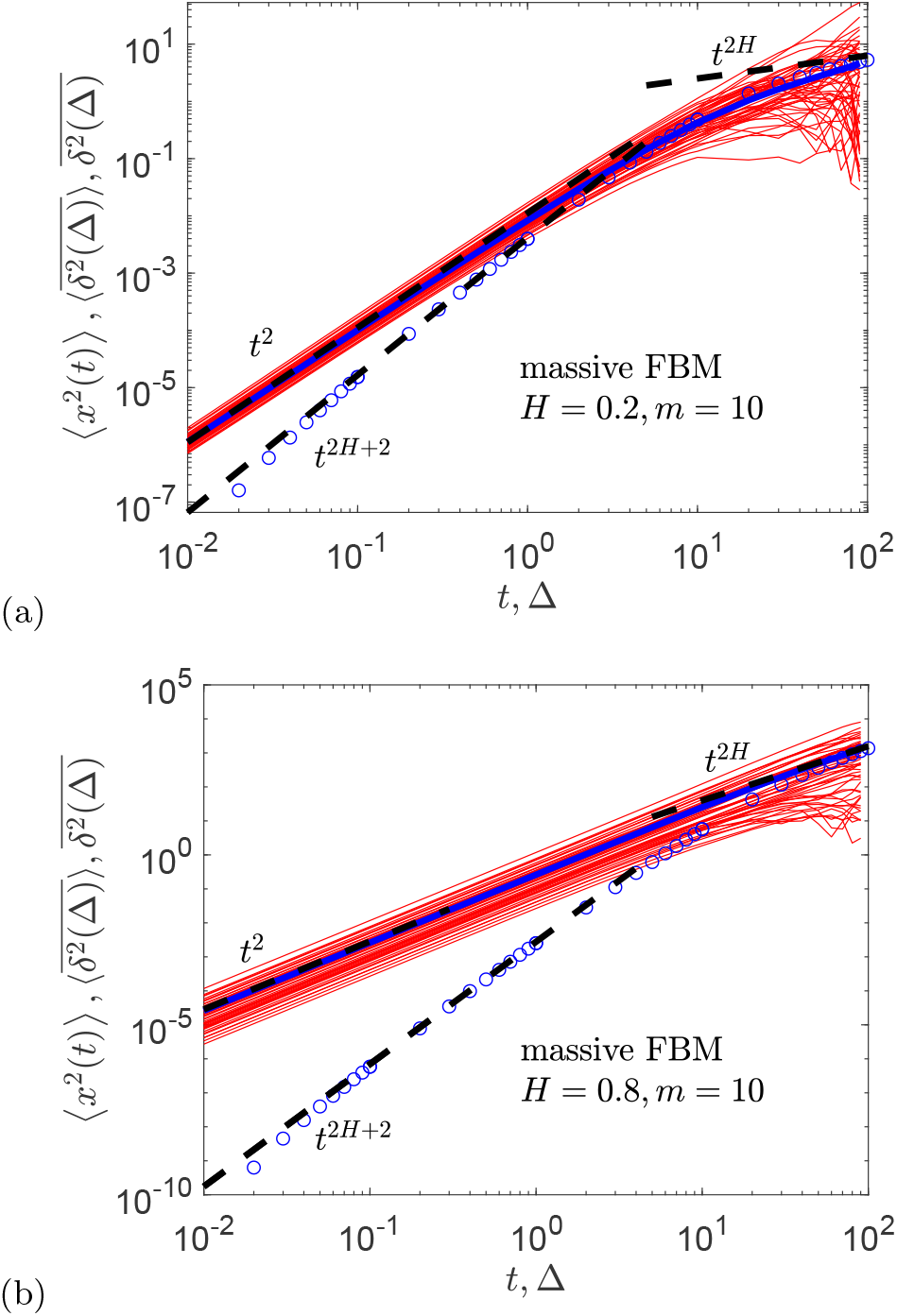
The same as in Fig. 1, for the same parameters and with the same asymptotes being shown, except for heavier particles, with *m* = 10.

**FIG. S2:**
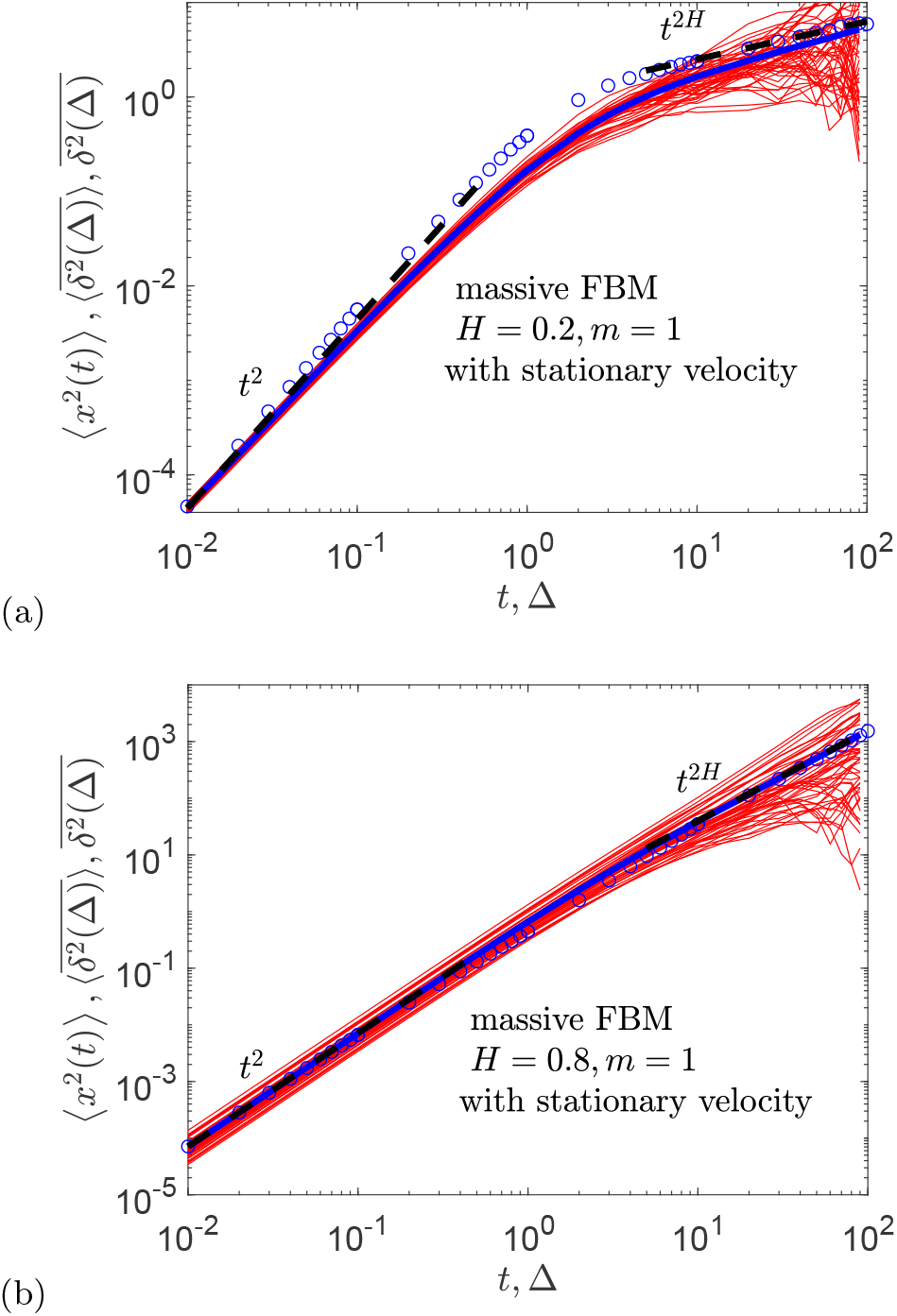
The same as in Fig. 1, for the same model parameters, except for the initial particle velocities taken from the distribution (A11).

**FIG. S3:**
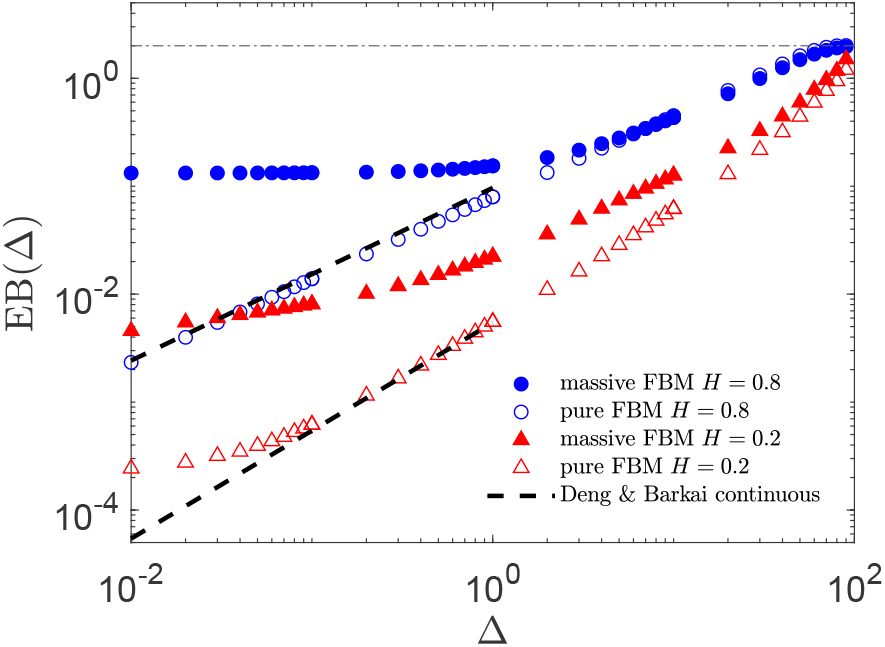
EB parameter (8) plotted versus lag time Δ for two values of the Hurst *H* exponent (see the legend). The theoretical continuous-time asymptote (19) by Deng-Barkai^7^ and the terminal EB value (23) are shown as the dashed and dot-dashed black lines, respectively. Parameters: *T* = 10^2^, *dt* = 10^−2^.

**FIG. S4:**
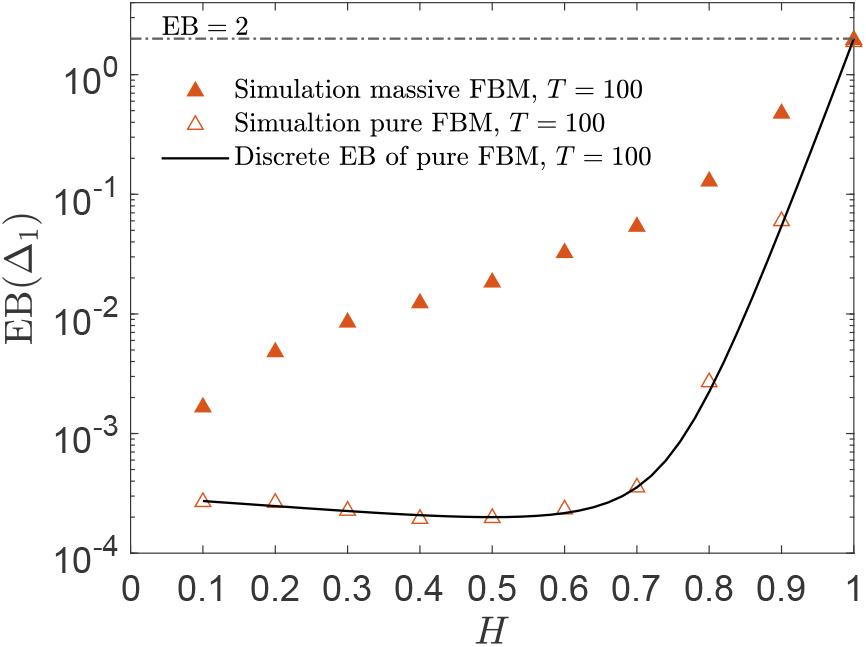
Variation of EB(Δ_1_) of overdamped FBM from the simulation data (as those in Fig. S3) versus the Hurst exponent *H*. The prediction of the discrete-time theory of EB for massless FBM^13,22^ given by (22) is also shown (see the legend). Parameters: *m* = 1, *T* = 10^2^, and Δ_1_ = *dt* = 10^−2^.

**FIG. S5:**
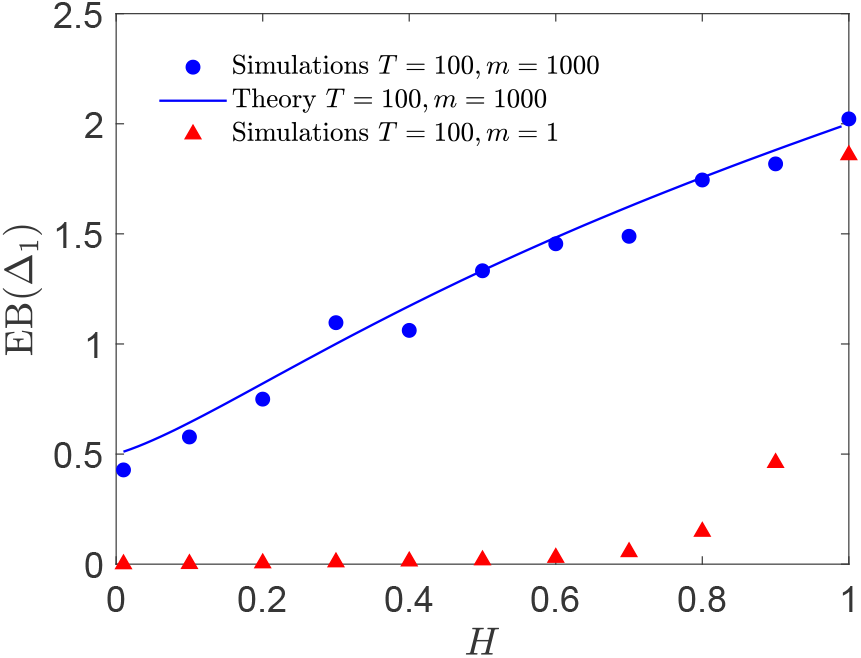
EB parameter of massive FBM in the limit *T* ≪ *m/γ* = *τ*_0_ (the blue symbols) and *T ≫ τ*_0_ (the red symbols, see the legend). The analytical short-trace or large-mass EB asymptote given by expression (A40) is the solid blue curve for the case *T* ≪ *τ*_0_.

**FIG. S6:**
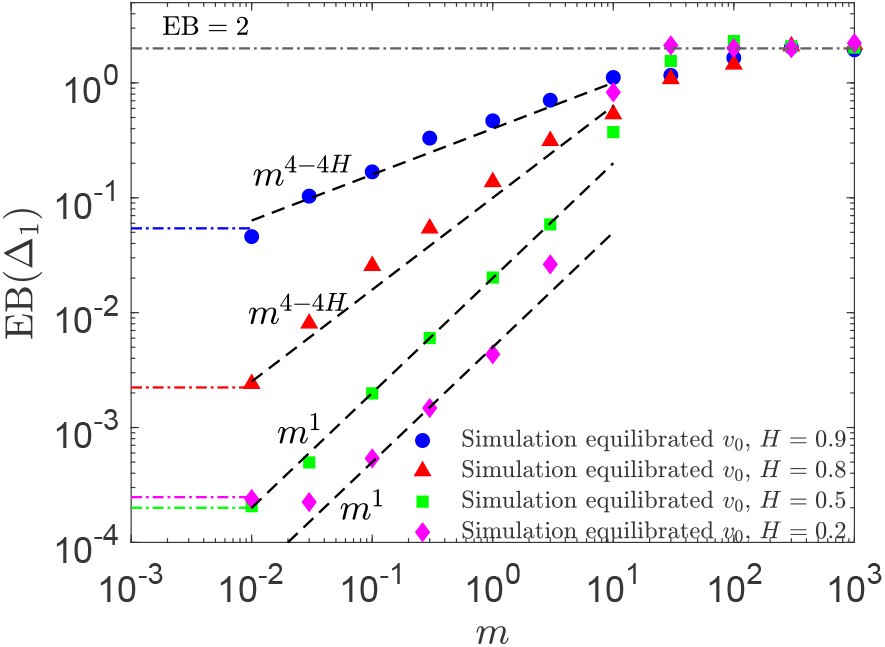
The same as in Fig. 3, for the same parameters and with scaling relations (24) shown, but computed for initial velocities distributed as in the stationary state, see Eq. (A11). The large-mass-asymptote (A43) is the dot-dashed black line at EB=2.

**FIG. S7:**
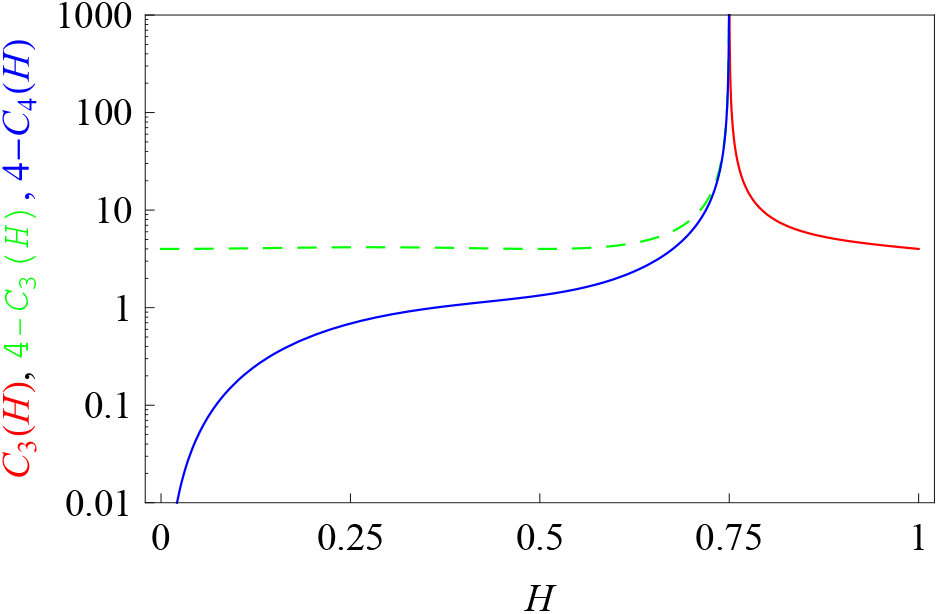
Prefactors in expressions (A48) and (A51), with a visible divergence at “critical” Hurst exponent *H* = 3*/*4.

